# Sensory uncertainty influences motor learning differently in blocked versus interleaved trial contexts when both feedforward and feedback processes are engaged

**DOI:** 10.1101/2023.11.28.569131

**Authors:** Matthew J. Crossley, Christopher L. Hewitson, David M. Kaplan

**Affiliations:** School of Psychological Sciences, Macquarie University, Sydney, Australia; Macquarie University Performance and Expertise Research Centre, Macquarie University, Sydney, Australia; Department of Psychology, Yale University, New Haven, USA; Wu Tsai Institute, Yale University, New Haven, USA

## Abstract

Theories of human motor learning commonly assume that the degree to which movement plans are adjusted in response to movement errors scales with the precision of sensory feedback received regarding their success. However, support for such error-scaling models has mainly come from experiments that limit the amount of correction that can occur within an ongoing movement. In contrast, we have recently shown that when this restriction is relaxed, and both within-movement and between-movement corrections co-occur, movement plans undergo large and abrupt changes that are strongly correlated with the degree of sensory uncertainty present on the previous trial and are insensitive to the magnitude and direction of the experienced movement error. Here, we show that the presence of these abrupt and error-insensitive changes can only be reliably detected when different levels of sensory precision are interleaved pseudo randomly on a trial-by-trial basis. These results augment our earlier findings and suggest that the co-occurrence of within-movement and between-movement corrections is not the only important aspect of our earlier study that challenged the error-scaling models of motor learning under uncertainty.

**Author summary:** A large body of literature shows that sensory uncertainty inversely scales the degree of error-driven corrections made to motor plans from one trial to the next. However, by limiting sensory feedback to the endpoint of movements, these studies prevent corrections from taking place during the movement. We have recently shown that when such corrections are promoted, sensory uncertainty punctuates between-trial movement corrections with abrupt changes that closely track the degree of sensory uncertainty but are insensitive to the magnitude and direction of movement error. Here, we show that this result requires different levels of sensory uncertainty to be mixed on a trial-by-trial basis. This carries important implications for how previous studies of motor learning under uncertainty are interpreted, and what future studies will likely constitute progress for the field.

## Introduction

Standard models of motor learning assume that the rate of feedforward adaptation – i.e., the process of adjusting motor plans in response to movement errors – is (1) a linear function of error size [1–7] and (2) scales with the precision (or equivalently inversely scales with the uncertainty) of the sensory signals relaying movement outcomes [8–12]. Thus, according to these *error-scaling* models the motor system will adapt more for large errors than for small errors, and more for highly certain sensory feedback than it will for highly uncertain sensory feedback.

Most studies that support *error-scaling* models of motor learning under uncertainty have used designs that deliberately limit the possibility of corrections being made within-movement. This choice reflects the desire to observe motor planning free from the potentially confounding influence of online motor control. However, we have recently reported that when within-movement error corrections are promoted, sensorimotor adaptation does not appear consistent with *error-scaling* models [13]. In particular, we showed that under these conditions movement plans undergo large and abrupt changes that are strongly correlated with the degree of sensory uncertainty present on the previous trial but are insensitive to the magnitude and direction of the experienced movement error.

Another aspect of our design that differed from the existing literature is that we exposed participants to different levels of sensory uncertainty interleaved pseudorandomly across trials, whereas most previous research used blocked designs in which one level of sensory uncertainty was presented for many consecutive trials [12, 14–16]. In one exception to this trend, [11] showed that error-scaling models hold in an interleaved design that limits within-movement correction. We reasoned on the basis of this study that the principle driver of our results was most likely to be that ours promoted within-movement corrections. However, it is possible that both the promotion of within-movement error correction and also trial-by-trial variation in sensory uncertainty play important roles in determining the adequacy of *error-scaling* models. We seek to empirically investigate that possibility here.

## Results

The aim of our experiments was to examine how sensory uncertainty influences motor learning when both within-movement and between-movement corrections co-occur and when different levels of sensory uncertainty are applied pseudorandomly across trials versus when they occur in a large block of trials. Participants made planar reaching movements using visual feedback about their hand position provided immediately before movement onset, at movement midpoint, and at movement endpoint. On low sensory uncertainty trials (*σ*_*L*_), this feedback took the form of a single cursor, and on high sensory uncertainty trials (*σ*_*H*_), this feedback took the form of a cloud of cursors (see the “Methods” section for details). We quantify our behavioral results using traditional statistical methods (see the “Statistical modeling” section for details) as well as by fitting a state-space model that make explicit assumptions about how motor commands are planned, executed, and updated over time, as well as how these processes are modulated by sensory uncertainty (see section “State-space modeling” for details.)

Fig 1 shows group-averaged initial movement vectors separately for each uncertainty level for both *blocked* and *interleaved* conditions. Blue indicates a trial in which low uncertainty feedback (*σ*_*L*_) was encountered on the *previous* trial. Orange indicates high uncertainty feedback (*σ*_*H*_) was encountered on the *previous* trial. The most striking aspect of these results is how much greater the initial movement vectors are for low uncertainty trials in the *interleaved* condition (dashed blue line) as compared to in the *blocked condition*. Note that this occurs despite the fact that there are far fewer low uncertainty trials in the *interleaved* condition than there are in the *blocked* condition.

**Fig 1.**
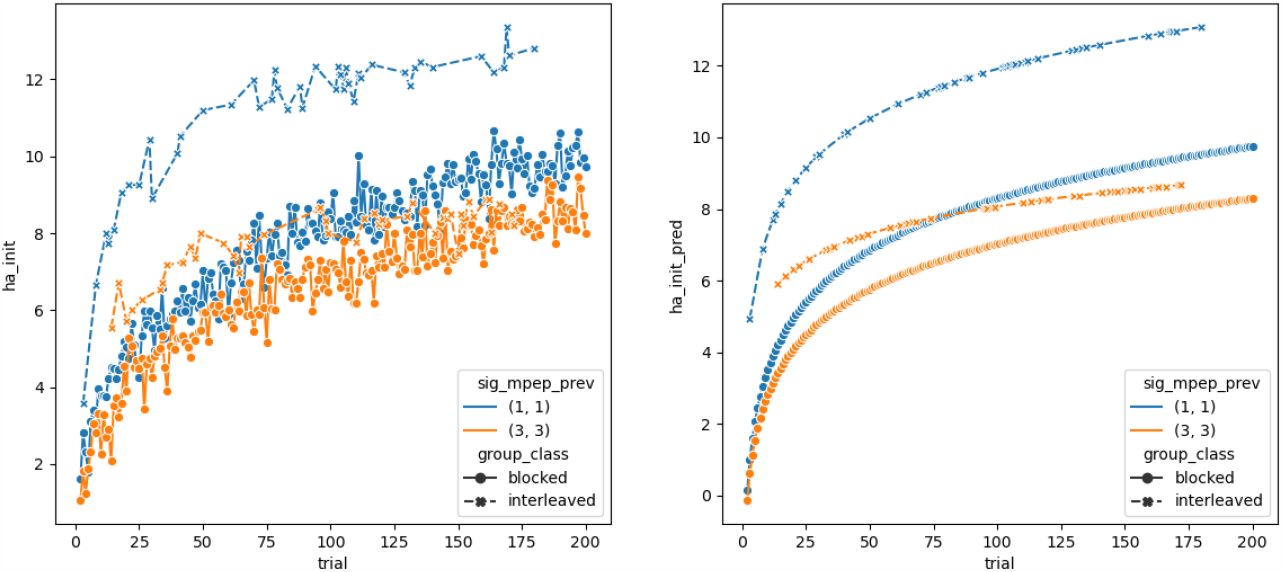
Feedforward adaptation results. (Left) Mean initial movement vector per trial across all participants. (Right) Initial movement vector predictions from the regression model. The solid line connected by filled circles are the *blocked* conditions and the dashed line connected by x’s are the *interleaved* condition.

It is also clear from this figure that greater sensory uncertainty leads to slower learning than lower sensory uncertainty in the *blocked* conditions (i.e., the solid blue line is consistently above the solid orange line). While it is tempting to say the same for the *interleaved* condition (i.e., the dashed blue line is also above the dashed orange line), it would be inaccurate to do so. This is because this condition is within-subject, so the large gap between blue and orange curves actually indicates large and abrupt changes in initial movement vectors oscillating between them.

Initial movement vectors over all trials and sensory uncertainty levels were greater in the *interleaved* condition than they were in the *interleaved* condition (Table 1 row 1). *log*(*Trial*) significantly predicted initial movement vectors (Table 1, row 9), indicating that initial movement vectors tend to increase over the course of the adaptation phase (Table 1 row 4). The interaction between condition and *log*(*trial*) was significant indicating that initial movement vectors increased across trials more slowly in the *blocked* conditions than in the *interleaved* condition (Table 1 row 5). The interaction between sensory uncertainty level and *log*(*trial*) was significant indicating that initial movement vectors increased across trials more quickly for low sensory uncertainty levels than it did for low uncertainty levels (Table 1 row 6). The three-way interaction between condition, sensory uncertainty level, and trial was significant indicating that initial movement vectors increased across trials more quickly for low sensory uncertainty trials in the interleaved condition compared to in the blocked conditions (Table 1 row 7).

**Table 1.**
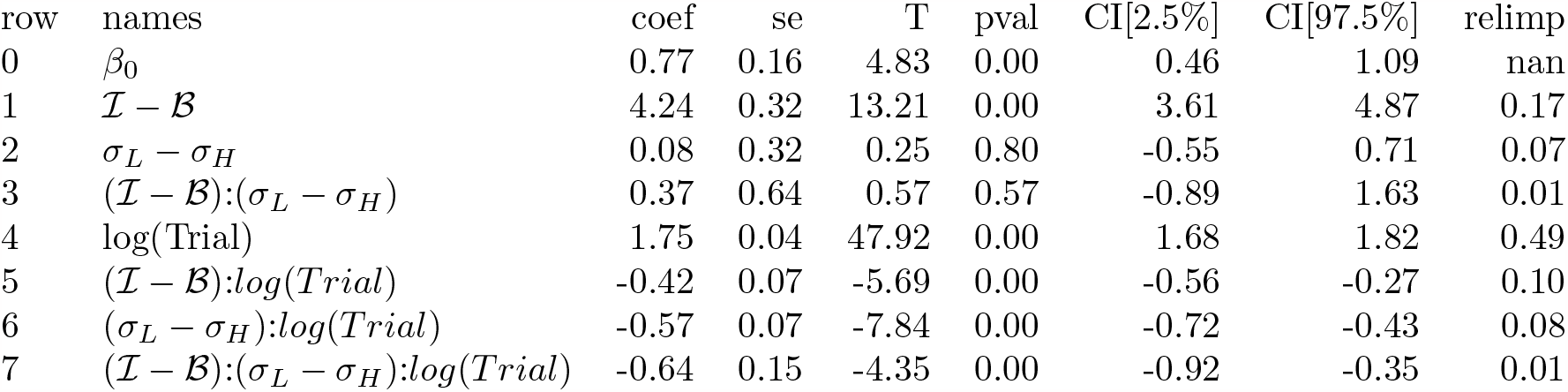
Experiment 1 regression results for predicting initial movement vector from error and sensory uncertainty terms. *ℐ* stands for *Interleaved* and *ℬ* stands for *Blocked*. The *coef* column contains *β* coefficients, the *se* column contains standard errors of these coefficients, the *T* column contains corresponding t-statistic, the *pval* column contains corresponding p-values, the *CI[2*.*5%]* and *CI[97*.*5%]* columns give the 95% confidence interval, and the *relimp* column gives the corresponding relative importance.

| row | names | coef | se | T | pval | CI[2.5%] | CI[97.5%] | relimp |
| --- | --- | --- | --- | --- | --- | --- | --- | --- |
| 0 | $\beta_0$ | 0.77 | 0.16 | 4.83 | 0.00 | 0.46 | 1.09 | nan |
| 1 | $\mathcal{I} - \mathcal{B}$ | 4.24 | 0.32 | 13.21 | 0.00 | 3.61 | 4.87 | 0.17 |
| 2 | $\sigma_L - \sigma_H$ | 0.08 | 0.32 | 0.25 | 0.80 | -0.55 | 0.71 | 0.07 |
| 3 | $(\mathcal{I} - \mathcal{B}):(\sigma_L - \sigma_H)$ | 0.37 | 0.64 | 0.57 | 0.57 | -0.89 | 1.63 | 0.01 |
| 4 | $\log(\text{Trial})$ | 1.75 | 0.04 | 47.92 | 0.00 | 1.68 | 1.82 | 0.49 |
| 5 | $(\mathcal{I} - \mathcal{B}): \log(\text{Trial})$ | -0.42 | 0.07 | -5.69 | 0.00 | -0.56 | -0.27 | 0.10 |
| 6 | $(\sigma_L - \sigma_H): \log(\text{Trial})$ | -0.57 | 0.07 | -7.84 | 0.00 | -0.72 | -0.43 | 0.08 |
| 7 | $(\mathcal{I} - \mathcal{B}):(\sigma_L - \sigma_H): \log(\text{Trial})$ | -0.64 | 0.15 | -4.35 | 0.00 | -0.92 | -0.35 | 0.01 |

The relative importance of the *log*(*Trial*) term in the regression model was 0.49, indicating that the slow envelope of the adaptation curve is well captured by a simple logarithmic function. The combined relative importance of the difference in conditions *ℐ* − *ℬ* and the interaction of this term with *log*(*Trial*) was 0.27, indicating that there is a substantial difference between conditions. The combined relative importance of the difference in uncertainty level *σ*_*L*_ − *σ*_*H*_ and the interaction of this term with *log*(*Trial*) was 0.17, indicating that there is a substantial difference between uncertainty levels. Overall, this regression analysis revealed a clear effect of sensory uncertainty and a clear effect of blocked versus interleaved condition.

We have previously shown that the pattern of results in the *interleaved* condition is difficult to reconcile with traditional error-scaling accounts of how sensory uncertainty influences implicit motor adaptation [13]. This is due to the large and abrupt trial-by-trial fluctuations in hand angle seen in this condition that seem almost entirely dictated by the sensory uncertainty experienced on the previous trial, and have little to do with the error experienced on that same trial. On the other hand, the pattern of results in the *blocked* conditions appear quite consistent with traditional error-scaling models. That is, adaptation under low sensory uncertainty seems to occur slightly more quickly and to a slightly greater extent than adaptation under high sensory uncertainty. To formalize this observation, we fit a single-state state-space model to the *blocked* conditions (see the “State-space modeling” section for details).

The results of these fits are shown in Fig. 2. The first important take-away from this figure is that a single-state model provides a compelling fit to the *blocked* data (as shown in the left panel of Fig. 2). This stands in sharp contrast to the *interleaved* condition, for which single-state models perform very poorly (see [13] and [17] for an illustration and discussion of this fact). The second important take-away from this figure is that the difference in sensory uncertainty between the two *blocked* conditions is captured in the model by differences in the learning rate parameter (Fig. 2 middle; *t*(19) = 3.03, *p <* 0.01, *d* = 0.96) and not by differences in the retention parameter (Fig. 2 right; *t*(19) = −1.77, *p* = 0.09, *d* = 0.46). Therefore, these modeling results are consistent with the predictions of traditional *error-scaling* models.

**Fig 2.**
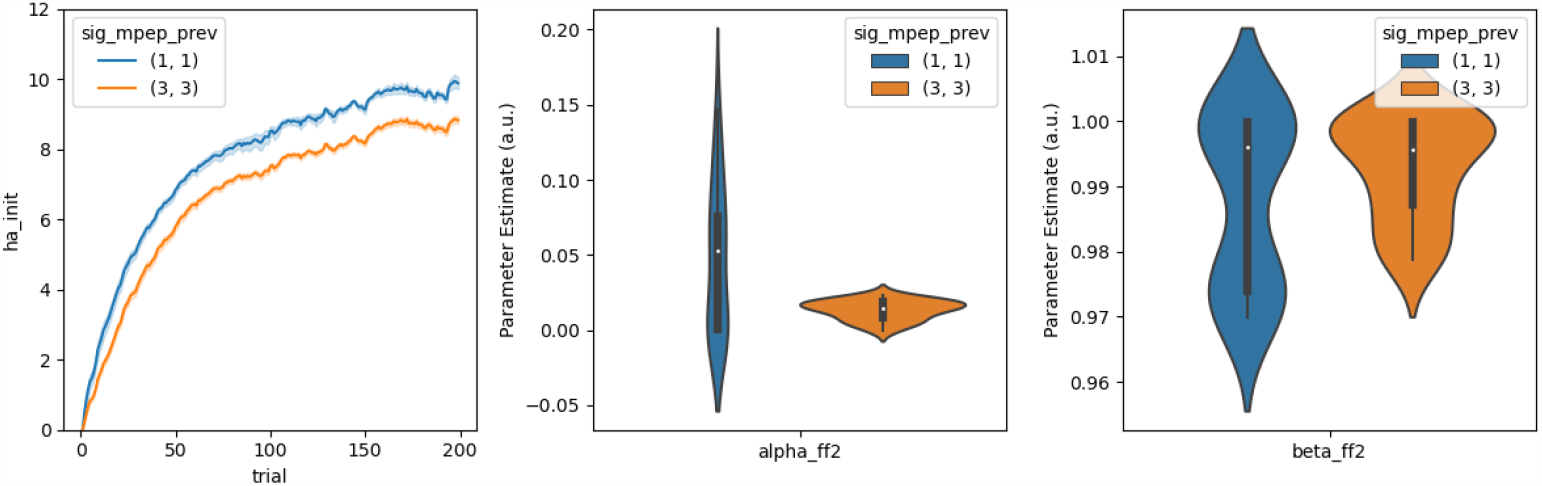
State-sapce model results. (Left) Mean initial movement vector per trial predicted from a single-state state-space model. (Middle) Best-fitting learning rate parameters (*α*). (Right) Best-fitting retention parameters (*β*).

## Discussion

The current study builds on our first examination of how sensory uncertainty influences motor learning in a context where within-movement corrections are promoted [13]. There we showed that transitions between different levels of sensory uncertainty on a trial-by-trial interleaved basis produces by large, abrupt, and frequent changes in initial movement vectors that are insensitive to the magnitude and direction of sensed movement errors. This pattern of behavior presents an obvious challenge to standard *error-scaling* models of how sensory uncertainty influences motor adaptation. Here, we show that this result is only evident when sensory uncertainty varies from trial to trial. It does not persist when sensory uncertainty is held is held constant across relatively large blocks of trials. This carries important implications for how previous studies of motor learning under uncertainty are interpreted, including ours, and what future studies will likely facilitate progress for the field.

Standard *error-scaling* models of motor learning assume that the amount of adaptation that will occur in response to a movement error is (1) proportional to the magnitude of that error but goes in the opposite direction [1–7], and (2) is proportional to the sensory precision (or equivalently inversely proportional to the sensory uncertainty) in signal conveying the movement error [8–12]. Our earlier results – e.g., those in the *interleaved* condition – challenged this characterization [13]. In particular, while the data in the *interleaved* condition is characterized by a slow envelope of error reduction consistent with *error-scaling* (i.e., both the blue and orange dashed lines in Fig. 1 gradually increase), it is also – and perhaps most prominently – characterized by the large difference in initial movement vectors between trials following low versus trials following high sensory uncertainty. Since these trials are interleaved, this means that participants reliably change their hand angle following high uncertainty trials in a direction that causes them to experience greater error than would be experienced if they simply ignored the feedback altogether. Standard *error-scaling* models posit that high sensory uncertainty should reduce adaptation rate, but they do not in any way suggest that adaptation should be reversed following high sensory uncertainty.

How might we have observed such radically different behavioral responses to sensory uncertainty than has previously been observed in over 20 years of investigation? We previously suggested that the critical aspect of our experiment was that it promoted within-movement error correction whereas most other studies deliberately suppressed it (see the “Key paradigm differences” discussion section in [13] for details). Here we considered another important design difference. Most studies of how sensory uncertainty influences motor learning have used blocked designs [12, 14–16], and these studies have all reported results consistent with classic *error-scaling* models. Only two previous studies have used interleaved designs. Körding and Wolpert (2004) [18] used a design virtually identical to those of our *interleaved* condition. They examined movements in the presence of within-movement and between-movement corrections and also with interleaved uncertainty conditions. And yet theirs is the seminal study that helped establish error-scaling models in the first place. However, since they did not report early adaptation trials, a direct comparison with our results impossible. Wei and Körding (2010) [11] also used a trial-interleaved design – though this study did not promote within-movement corrections as we have done here – and also found support for an error-scaling model.

The present findings shed new light on our previous results. In particular, the pattern of behavior observed in the *blocked* conditions appears highly consistent with classic error-scaling accounts of how sensory uncertainty influences motor learning. Thus, given our present data, it seems that the error-scaling model can account for motor learning under uncertainty when within-movement and between-movement corrections are promoted, provided that different uncertainty conditions are not interleaved across trials.

Where the *error-scaling* model fails to capture the pattern of behavior seen in the *interleaved* condition, an *aiming model* assuming that different levels of sensory uncertainty trigger different aiming strategies succeeds [13, 17]. The aiming explanation not only provides a compelling model fit to the data, it also has the appeal of avoiding the need to explain what would otherwise be a bizarre pattern of adaptation. However, as we articulated in our earlier work [13], it precipitates some puzzles of its own. In particular, it is unclear why a successful aiming strategy would subsequently be abandoned in favor of a less successful aiming strategy on subsequent trials, only to return to the original aiming strategy just a few trials later (i.e., as is implied by the stratification of initial movement vectors across different levels of sensory uncertainty apparent in our data). The present results add to this puzzle. In particular, when viewed through the lens of explicit aiming, they suggest that more extreme aiming occurs, especially for the low uncertainty condition (*σ*_*L*_), when different levels of sensory uncertainty are interleaved versus when they are blocked. Since no theory of aiming makes this prediction a priori, the present results cast further suspicion on the possibility that aiming is at the heart of the interleaved results.

It may also be useful to interpret our results through the lens of contextual inference [17]. According to several influential theories, motor learning is exquisitely context-specific [19–21]. Though varied in their implementation and specific assumptions, these theories roughly assume that the brain maintains many internal models that learn context-appropriate sensorimotor mappings. From this perspective, it is possible that different sensory uncertainty conditions engage different context-specific internal models. However, this explanation also precipitates its own set of puzzles. In particular, it implies that the learning rate assigned to the low uncertainty context is greater when trial types are interleaved versus when they are blocked. None of the context models cited above make this type of prediction a priori. Consequently, more experimental and theoretical work is required to resolve these important questions.

## Conclusion

Real-world human movement involves both within-movement and between-movement error correction, and with similar inevitability occurs within the context of sensory uncertainty. The present study builds on our recent paper [13], which was the first to examine how sensory uncertainty influences motor learning when within-movement error corrections is promoted. We have previously shown that classic *error-scaling* models of how sensory uncertainty influences motor learning do not offer viable or parsimonious account of human reaching behavior when observed in an experimental context that promotes within-movement error correction [13]. The results of the present study suggest that the adequacy of *error-scaling* models depends on the confluence of promoting within-movement corrections with the interleaving of differing levels of sensory uncertainty across trials.

## Materials and methods

### Participants

A total of 60 naive participants (20 per condition; aged 18-32; 28 male; 32 female) with normal or corrected to normal vision and no history of motor impairments participated in the experimental study. All participants gave written informed consent before the experiment and were either paid and recruited from the Macquarie University Cognitive Science Participant Register or were Macquarie University undergraduates participating for course credit.

All experimental protocols were approved by the Macquarie University Human Research Ethics Committee (protocol number: 52020339922086). Participants were randomly assigned to one of three conditions (n=20 per experiment). Sample sizes were consistent with field-standard conventions for visuomotor adaptation experiments [22–24]. The data from the *interleaved* condition has been previously reported in Experiment 2 of our recent publication [13].

### Experimental Apparatus

A unimanual KINARM endpoint robot (BKIN Technologies, Kingston, Ontario, Canada) was utilized in the experiments for motion tracking and stimulus presentation. The KINARM has a single graspable manipulandum that permits unrestricted 2D arm movement in the horizontal plane. A projection-mirror system enables presentation of visual stimuli that appear in this same plane. Participants received visual feedback about their hand position via a cursor (solid white circle, 2.5 mm diameter) controlled in real-time by moving the manipulandum. Mirror placement and an opaque apron attached just below the subject’s chin ensured that visual feedback from the real hand was not available for the duration of the experiment.

### General Experimental Procedure

Participants performed reaches with their dominant (right) hand – as assessed by the Edinburgh handedness inventory (¿40 equates to right-hand dominant) – from a starting position located at the center of the workspace (solid red circle, 0.5cm in diameter) to a single reach target (solid green circle, 0.5 cm in diameter) located straight ahead (0° in the frontal plane) at a distance of 10 cm. When participants moved the cursor within the boundary of the start target its color changed from red to green and the reach target appeared, indicating the start of a trial. Participants were free to reach at any time after the start target color changed.

Participants in all conditions first completed a 20 trial baseline phase during which veridical online feedback was provided. Immediately following baseline, an adaptation phase was completed (180 trials in the *interleaved* condition and 200 trials in the *blocked* conditions) during which the cursor feedback was extinguished once the cursor exited the start target, but briefly reappeared (100ms duration) at movement midpoint and at movement endpoint. These brief bouts of visual feedback were rotated counterclockwise (to the left) of the true hand position by an amount drawn at random on each trial from a Gaussian distribution with a fixed mean of 12° and standard deviation of 4°. This random sequence of rotations was identical for each participant within a given condition – and was identical in each *blocked* condition – but differed between the *blocked* and *interleaved* conditions.

In the *blocked* condition, the brief bouts of visual feedback presented at movement midpoint and endpoint were displayed with either low visual uncertainty (*σ*_*L*_) or high visual uncertainty (*σ*_*H*_). In the *interleaved* condition these bouts of visual feedback were displayed at one of four visual uncertainty levels (*σ*_*L*_, *σ*_*M*_, *σ*_*H*_, *σ*_*∞*_). The sequence of uncertainty levels across trials was random – subject to the constraint that 45 trials of each type would be sampled in total – but was identical for each participant. Here, we only report *σ*_*L*_ and *σ*_*H*_ trials in order to facilitate comparison to the *blocked* conditions. For a full analysis of the *interleaved* data, please consult our earlier work [13].

In the low uncertainty condition (*σ*_*L*_), feedback was a single white circle (0.5 cm in diameter; 5.73° arc-angle at midpoint, 2.86° at endpoint), identical to the initial cursor. In the moderate uncertainty condition (*σ*_*M*_), feedback was one of 10 randomly generated point clouds comprised of 50 small translucent white circles (0.1 cm in diameter) distributed as a two-dimensional Gaussian with a standard deviation of 0.5cm (5.73° arc-angle at midpoint, 2.86° at endpoint), and a mean centered over the true (perturbed) cursor position on the current trial. In the high uncertainty condition (*σ*_*H*_), everything was the same as the moderate uncertainty condition (*σ*_*M*_) except that the point clouds had a SD of 1 cm 11.47° arc-angle at midpoint, 5.73° at endpoint). In the unlimited uncertainty condition (*σ*_*∞*_), no feedback was provided at all.

Participants were instructed to use cursor feedback to guide their reaches whenever it was available, and to move their hand straight through the target as accurately as possible. The maximum allowable time to complete a reach was 1000 ms. If participants did not move past the lower bound of the end target radius (9.5cm) the trial would time out and restart. If reaches exceeded the time limit or did not cross the lower bound of the target, the trial was repeated. To help guide the participant’s hand back to the starting position, a green ring centered over the starting position appeared with a radius equal to the distance between the hand and starting position. Once the participant’s hand was 1 cm from the starting position, the ring was removed and veridical cursor feedback was reinstated. Immediately following the adaptation phase, participants in all conditions completed a 100 trial washout phase during which no cursor feedback was provided. For simplicity and brevity, we do not report the washout data here, but will do so in an upcoming paper. There were no breaks between phases, and transitions between phases were not explicitly signaled to participants in any way.

### Data Analysis

Movement kinematics including hand position and velocity were recorded for all trials using BKIN’s Dexterit-E experimental control and data acquisition software (BKIN Technologies). Data was recorded at 200 Hz and logged in Dexterit-E. Custom scripts for data processing were written in MATLAB (R2013a). Data analysis and model fitting was done in Python (3.7.3) using the numpy (1.19.2) [25], SciPy (1.4.1) [26], pandas (1.1.3) [27], matplotlib (3.3.2) [28], and pingouin (0.3.11) [29] libraries.

A combined spatial- and velocity-based criterion was used to determine movement onset, movement offset, and corresponding reach endpoints [30, 31]. Movement onset was defined as the first point in time at which the movement exceeded 5% of peak velocity after leaving the starting position. Movement offset was similarly defined as the first point in time at which the movement dropped below 5% of peak velocity after a minimum reach of 9.5 cm from the starting position in any radial direction, and reach endpoint was defined as the (*x, y*) coordinate at movement offset. We used initial movement vector (IMV) – i.e., the angular difference between the target location and the movement vector at movement onset – as our estimate of feedforward adaptation.

### Statistical modeling

We fit a regression model to quantify the effect of condition (*interleaved* vs *blocked*) and sensory uncertainty (*σ*_*L*_ vs *σ*_*H*_) on feedforward adaption. We treated initial movement vector as the observed variable. Predictor variables were trial, condition, the sensory uncertainty experienced at midpoint / endpoint *on the previous trial*, and all interactions between the condition term, the sensory uncertainty term, and the trial term. We used backward difference coding to enter both condition and sensory uncertainty into this regression model. According to this coding scheme, the performance with sensory uncertainty at one level is compared with performance when sensory uncertainty is at the previous level. Thus, the regression models contains beta coefficients that capture the difference between the *interleaved* and the *blocked* condition as well as beta coefficients that capture the difference between low sensory uncertainty and high sensory uncertainty. This also applies to the interaction terms.

In all models for which trial number was taken to be a predictor, trial number was transformed using base-ten logarithm. This has the effect of turning our non-linear adaptation curves into straight lines, and thereby makes linear regression a more appropriate analysis tool for our research question. The models were fit to the group averaged data using ordinary least squares to obtain best-fitting parameter estimates. We also report the relative importance of each regressor following the methods developed in [32].

### State-space modeling

We fit a simple discrete-time linear dynamical system – i.e., a state-space model [5] – to the *blocked* conditions. This model includes one internal state variable to control feedforward movement plans (i.e., this state variable controls initial movement vectors), and one internal state variable to control within-movement error correction (i.e., this state variable influences how much initial movement vectors are altered in response to midpoint feedback). Since within-movement error corrections are not the focus of this paper, we leave the details of this process to our earlier work [13]. Moreover, updates to the internal state variable governing feedforward motor plans in this model depends on movement error sensed both at midpoint and also at endpoint. However, since the uncertainty of the sensory feedback delivered in all experiments considered here was matched at both time points, this dependency makes little difference, so we omit it from the following model description. Please see our earlier work for full details [13].

The feedforward component of this model assumed that an internal state variable *x* maps desired motor goals to motor plans *y*, and that *x* is updated on a trial-by-trial basis in response to movement error:

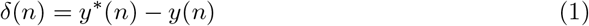

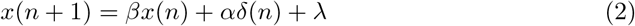

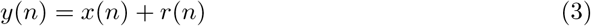

Here, *n* is the current trial, *δ* is the movement error, *y*^*\**^ is the desired output, *y* is the actual motor output, *r* is the imposed rotation, *x* is the state of the system, *β* is a retention rate that describes how much is retained from the value of the state at the previous trial, *α* is a learning rate that describes how quickly states are updated in response to errors, and *λ* is a constant bias.

We obtained best-fitting parameter estimates on a per subject basis by minimizing the sum of squared error difference between the observed and predicted midpoint and endpoint hand angles. We found the best-fitting parameter values using the differential evolution optimization [33] method implemented in *SciPy* [26].

## Code Accessibility

Data and analysis code can be accessed at: https://github.com/crossley/sensory_uncertainty_fffb_blocked_vs_interleaved

